# Whisper: Read sorting allows robust mapping of sequencing data

**DOI:** 10.1101/240358

**Authors:** Sebastian Deorowicz, Agnieszka Debudaj-Grabysz, Adam Gudyś, Szymon Grabowski

## Abstract

**Motivation:** Mapping reads to a reference genome is often the first step in a sequencing data analysis pipeline. Mistakes made at this computationally challenging stage cannot be recovered easily.

**Results:** We present Whisper, an accurate and high-performant mapping tool, based on the idea of sorting reads and then mapping them against suffix arrays for the reference genome and its reverse complement. Employing task and data parallelism as well as storing temporary data on disk result in superior time efficiency at reasonable memory requirements. Whisper excels at large NGS read collections, in particular Illumina reads with typical WGS coverage. The experiments with real data indicate that our solution works in about 15% of the time needed by the well-known Bowtie2 and BWA-MEM tools at a comparable accuracy (validated in variant calling pipeline).

**Availability:** Whisper is available for free from https://github.com/refresh-bio/Whisper or http://sun.aei.polsl.pl/REFRESH/Whisper/

**Supplementary information:** Supplementary data are available at publisher Web site.

## 1 Introduction

Mapping high-throughtput sequencing (HTS) short reads onto a reference sequence (also called read alignment) is nowadays both an industrial process and traditionally an active research topic, with over 80 published tools so far (see (Fonseca *et al*., 2012) and http://wwwdev.ebi.ac.uk/fg/hts_mappers/ for comprehensive, yet incomplete lists). Clearly, this is not the end of the road as each year new proposals appear, competing in mapping quality, functionalities, and—last but not least—aligning speed and memory requirements. Not all these algorithms target DNA; there are also several specialized RNA, miRNA, and bisulfite mappers. In this work we focus on DNA mapping.

The problem is not simple, because exact matching of reads onto the reference genome is of little value. Due to sequencing errors and genomic variations, most reads can be aligned to the reference sequence only approximately. For these reasons, the mapping algorithm should be based on approximate string matching, with tolerance for several mismatches and indels. It is also noteworthy that not all reads occur uniquely in a reference genome. Therefore, returning only one match (even with the minimal possible error) is insufficient in some applications, e.g., CNV calling.

Mapping reads to a given reference genome sequence is often the first step in sequencing data analysis and mistakes made at this stage cannot be easily recovered later. On the other hand, producing billions of bases daily as a routine job of modern sequencers (for example, Illumina HiSeq 4000 can generate as much as 400 Gb a day) makes the alignment task challenging not only from the mentioned quality point, but also performance-wise. It is well-known that the genomic database growth outpaces the famous Moore’s law for computing hardware (Kahn, 2011; Deorowicz and Grabowski, 2013), which means that the only hope for overcoming this flood of data is finding smarter and more efficient algorithms.

On a high level, in each mapping tool two major design decisions have to be made upfront: what data structure is used for the reference sequence (e.g., a genome) and what is the essential algorithm of finding the best read alignment (or multiple sufficiently good alignments). The answer for the former question is, in most cases, a hash array on *k*-mers of the reference genome or a compact full-text index (especially the FM-index (Ferragina and Manzini, 2000; Lam *et al*., 2009)).

Mappers based on a hash table, like MAQ (Li *et al*., 2008), SOAP (Li *et al*., 2008), and SHRiMP2 (David *et al*., 2011) generally follow the seed-and-extend approach (Li and Homer, 2010). According to this strategy, the query sequence is divided into *k*-mers and the positions (seeds) of each *k-*mer in another (reference) sequence are retrieved from a hash table. After that, the seeds are extended, joined and aligned using, e.g., the classic Smith–Waterman alignment algorithm. Using an FM-index (especially in a variant supporting bidirectional search (Luo *et al*., 2013)), also allows to locate quickly snippets of the query string. The well-known representatives of this strategy are BWA (Li and Durbin, 2009), Bowtie2 (Langmead and Salzberg, 2012), SOAP3 (Liu *et al*., 2012), and GEM (Marco-Sola *et al*., 2012).

Finding the best read alignment(s) is, however, related to the assumed similarity distance. In the majority of the second generation sequencing platforms (notably, Illumina) the vastly dominating kind of errors are mismatches (Hamming distance), but to compete with high-quality solutions it is becoming more and more important to handle also indels (edit distance). Note also that a non-negligible fraction of known variants, e.g., in a human genome, are indels, which is another reason to support this similarity model.

Kart (Lin and Hsu, 2017) is an efficient solution handling short as long as long reads. The algorithm employs both the FM-index (BWT array) and a hash table, and adopts a divide-and-conquer strategy to separate a read into easy and hard regions. The latter ones require gapped alignment and the final alignment is composed from independently aligned separate regions.

Most of the widely used mapping tools are *best-mappers*, which try to identify one or at most a few best mapping locations for each read (Langmead and Salzberg, 2012; Li and Durbin, 2009; Marco-Sola *et al*., 2012; Li, 2013). In some downstream applications (including ChIP-seq experiments, copy number variation calling and RNA-seq transcript abundance quantification) it is desirable to identify all relevant locations, which is the goal of *all-mappers* (Weese *et al*., 2012; Kim *et al*., 2010; Siragusa *et al*., 2013; Cheng *et al*., 2015).

Another way of classifying algorithms is the read processing regime. Most solutions map one read at a time, yet some of the exceptions are Masai (Siragusa *et al*., 2013) and TREQ-CG (Mahmud and Schliep, 2014). Masai jointly maps read prefixes using a Patricia trie (called a radix tree in the cited work). As long as a pair of reads has a common (exact) prefix, they are processed together, with a clear benefit for performance. TREQ-CG goes further, as it groups reads into clusters which are then represented with a single read being submitted to a mapping component. In fact, TREQ-CG is not a real read mapper; it is a clustering algorithm being a preprocessor for an arbitrary traditional mapper.

Time efficiency can be achieved not only with purely algorithmic means. Parallel processing can be found in many modern bioinformatic tools and read mappers are no exception. For example, BWT-based alignment can be implemented on massively parallel architectures like GPUs. Prominent examples are the tools SOAP3 (Liu *et al*., 2012) and SOAP3-dp (Luo *et al*., 2013), being about an order of magnitude faster than their CPU-based counterparts. The FPGA platform was also used for read mapping, with solutions involving a hash table (Olson *et al*., 2012) or an FM-index (Chen *et al*., 2013; Fernandez *et al*., 2015).

A problem related to the one considered in this work is mapping reads against multiple reference genomes. It has clear metagenomic applications, yet Schneeberger et al. (2009) argue that using multiple reference sequences for the same species should improve the mapping accuracy due to reducing the bias associated with a single genome. We agree this argument cannot be easily dismissed, yet the existing mappers to multiple genomes are rather immature.

For example, BWBBLE (Huang *et al*., 2013) needs more than 200 GB of memory to build a multi-genome for a collection of 1092 human genomes. GCSA (Sirén *et al*., 2014), in which the *pan-genome*, i.e., the reference genome and known variants of it, are represented with an extended BWT index of a graph, cannot be constructed in even 1 TB of RAM for a few “hard” human chromosomes (on the other hand, the resulting index is small). Both solutions need at least 10ms of time to find matches with up to 3 errors.

MuGI (Danek *et al*., 2014) is probably the only solution capable of finding all pattern occurrences in a collection of 1092 human genomes on a PC with 16 GB of RAM, searching for a pattern with 2 mismatches in well below 1 ms. We have to stress, however, that MuGI (like the other tools listed in the previous paragraphs) is an index rather than a full-fledged read mapper, since it handles mismatches only, does not support paired-end reads, and does not report alignment results to a SAM file.

A unique approach to read alignment was taken in two not well-known papers, presenting Slider (Malhis *et al*., 2009) and Syzygy (Konagurthu *et al*., 2010). Their strategy is called sort-and-join, as they lexicographically sort both the suffixes (or *k*-mers) from the reference string and the collection of reads, and then join both sorted sequences. Our work presented in this paper also follows this strategy. It has to be stressed, however, that although sort-and-join seems to be a good start, to obtain competitive mapping speed and quality we had to overcome major hurdles, e.g., related to significant amounts of repetitive genome areas.

This short overview supports our claim that industry-level multiple-genome read mappers are yet to come. There are also a number of theoretical works dedicated to indexing text with wildcard positions (Thachuk, 2013; Hon *et al*., 2013), where the wildcards represent SNPs, or the more general problem of indexing repetitive data with support for exact or approximate matching (Gagie *et al*., 2011; Jansson *et al*., 2014; Ferrada *et al*., 2014). None of them, however, can be considered a breakthrough, at least for bioinformatics, since none of them was demonstrated to run on multi-gigabyte genomic data (and in some of the cited papers no experimental results are given at all). The subject of indexing and searching genomic databases is also surveyed by Gagie and Puglisi (2015), with more focus on theoretical solutions.

## 2 Methods

### The general idea

As mentioned above, indexing the genome sequence is a canonical general approach to read mapping. One of the possible indexes to be used here is the well-known suffix array, in which all suffixes of the reference genomes are kept sorted, what allows to find *exact* matches with a binary search. This obvious strategy can be refined with backtracking to support approximate matching.

In this work, following Malhis *et al*. (2009) and Konagurthu *et al*. (2010), we propose to sort not only the reference sequence suffixes, but also the reads themselves. This has several benefits. Firstly, bulk processing of the reads is more efficient than taking them one by one to align against multiple distant locations in the reference sequence. The boost in speed can be explained by locality, and thus cache-friendliness, of bulk operations, where the successive sorted (and thus similar) reads are likely to be aligned to suffixes being close in their sorted order. This effect more than compensates the initial effort of sorting the reads. Secondly, the suffix array does not need to be wholly kept in the main memory, but instead may be read in successive portions from disk, matching the lexicographical range of the current portion of the sorted reads. In this way, we can enjoy the fast and convenient suffix array data structure without suffering from its space requirement, which is (at least) 4*n* bytes for sequences of length *n* < 4 Gb.

The whole algorithm can be divided into three phases: preprocessing, main processing, and postprocessing. We assume that two suffix arrays, *SA* for the reference genome and *SA*_rc_ for the reverse-complemented reference genome, are initially built and stored on disk. Still in the preprocessing, the reads’ DNA sequences are extracted from input FASTQ file(s) and written with some metadata on disk; the corresponding quality scores are ignored. In the main processing phase we look for possibly good alignments for all the reads, ignoring the fact that for paired-end (PE) data the expected alignments for a pair of reads are spatially correlated. In the postprocessing, for each pair of reads their “feasible” alternative alignments are merged, taking into account their genomic locations. If no such pair can be found, then the alignment for one of the reads is fixed (here many such candidate alignments may be tried one by one) and the other is scanned over the reference genome, in the close neighborhood with higher error threshold and clipping allowed.

Each of the phases is performed in parallel (with care for thread load balancing) and temporary data are stored on disk.

### Preprocessing

Multiple input FASTQ files are processed independently, yet the paired reads (for paired-end data) from “_1” and “_2” files are handled by twin threads, not to lose their association.

A vast majority of Illumina reads in an experiment are of the same length, yet it may happen that read lengths even from a single dataset vary. As handling reads of significantly different lengths would complicate the internals of Whisper, a simple rule of thumb is used: all the reads shorter than 90% of the length of the longest read (which is found during the proprocessing) are discarded. Practically, very few Illumina reads are removed in this way.

The DNA symbols from reads are filtered. All symbols other than `ACGT` are replaced with `N` (yet `N` symbols in the reference sequence have their own encoding, as `N`-to-`N` between a read and the reference at an aligned position should be considered as a mismatch). Each read is represented as a triple: (*i*) a unique 40-bit internal ID such that reads from a pair differ only in the least significant bit, (*ii*) DNA string, with non-standard symbols replaced by `N`, (*iii*) min_error, which is the error (i.e., the number of mismatches and indels) of the most successful read mapping found so far. This value is stored on a byte and is initialized to 255 (which is the maximum handled edit distance in our solution, yet the reads may be longer).

The reads are sent into multiple bins on disk. Bins are identified by their variable-length prefixes fulfilling the prefix property (no element in this set is a prefix of another). Those prefixes are established in a way that the distribution of the number of their occurrences in the reference is approximately uniform. As the maximum bin prefix length is set to 10, there are as many as 4^10^ different prefixes not containing symbol `N`, and their set is (essentially) iteratively reduced by merging the items down to 384 (by default).

### Main processing

We assume that all read alignments with up to *k* edit (Levenshtein) errors are to be found. Once the bins on disk are ready, we start the main processing phase, which can be divided into *k* + 2 major stages. We start with presenting the first *k* + 1 major stages (the last stage is different). The *i*th (0 ≤ *i* ≤ *k*) major stage is divided into *i* + 1 minor stages. In the *i*th major stage all matches with *exactly i* errors are found; the matches with less errors have already been found in the previous major stages. Reads, once matched, are no longer processed in the further major stages. To this end, the current read is divided into *i* + 1 (approximately) equal disjoint segments of length *m*/(*i* + 1), where *m* is the length of the shortest accepted read. If the current read is longer than *m*, its last segment is *not* longer than previous ones, but simply the remaining symbols are not a part of any segment (yet they are not ignored for the approximate matching performed later); for technical reasons it is convenient if the segments at a single substage are of equal length.

At the start of each *j*th (0 ≤ *j* ≤ *i*) minor stage of *i*th major stage, which will be denoted as stage (*i*, *j*), the reads are sorted according to their substring of length about *m*/(*i* + 1) being precisely the *j*th segment. Note that, by the Dirichlet principle frequently used in approximate string matching, if a string matches a reference with at most *k* Levenshtein errors, then at least one of its *k* + 1 disjoint pieces must occur in the considered alignment in an exact form (Rivest, 1976). The idea of having several multiple stages (instead of one, looking for matches with up to *k* errors) is to speed up computations. Matches found in earlier stages (in which the matching segments are longer) help reduce the number of read-suffix pairs to verify in later stages.

Let us consider an arbitrary stage (*i, j*). The bins are read from disk one by one. Assume one particular bin to be processed. The reads from the bin are first lexicographically sorted. The corresponding segments of the suffix arrays *SA* and *SA*_rc_ are also read from disk. We scan successive reads. If the current read is identical to the previous one (it happens quite often in large real collections), we simply copy the mapping results from its predecessor. Otherwise, we check if its *j*th segment only is equal to the *j*th segment of the previous read. If they are different, a range of suffixes in *SA* (resp. *SA*_rc_) matching this segment needs to be found. For efficiency reasons, including reducing the number of cache misses, we do not scan *SA* (resp. *SA*_rc_) linearly, but jump by the number of suffixes roughly equal to the ratio of the total number of suffixes in the segment of *SA* (resp. *SA*_rc_) corresponding to the current bin to the number of reads in the current bin. (More precisely, this jump size is actually greater by a factor of about 1.4, which was established experimentally.) If after this interpolation step we are still before the first desired suffix, we skip the same number of reads, etc. Once we go too far, a binary search over the last considered range of suffixes is performed. If, however, the *j*th segment of the read is equal to the *j*th segment of the previous read, there is no need to look for the relevant range of suffixes as this is simply copied from the previous read.

As each suffix in the relevant range must be tested against the current read, we do not (explicitly) look for the last suffix in a range; rather, we iterate linearly over the suffixes until the matching range ends. For each tested suffix, we compute the Levenshtein distance between it and the current read. If the distance is not greater than *i*, the mapping is recorded (in memory, so far). The approximate matching with up to *i* differences (i.e., *i* Levenshtein errors) is performed with Myers’ bit-parallel algorithm (Myers, 1998), unless *i* is small (*i* ≤ 7), when a dynamic programming procedure restricted to the 2*i* +1 central diagonals is used.

The found mappings are written to an output file as a quadruple: (*i*) read’s internal ID (40 bits), (*ii*) match position in *SA* (resp. *SA*_rc_), (*iii*)a flag if the match was against *SA* or *SA*_rc_,(*iv*) the distance (i.e., number of errors). We set a limitation of up to 1024 stored mappings per read. In each stage (*i, j*), *j* < *i*, the already processed reads are distributed into new bins for the next minor stage, i.e., stage (*i*, *j* + 1). In the last minor stages (the case of *i* = *j*), however, new bins are created for stages (*i* + 1, 0).

There are two practically important optimizations in the described procedure. One is a filter allowing not to compute the (relatively costly) Levenshtein distance between a read and a suffix, if a quick check tells that the distance exceeds the limit (which is *i* in the *i*th major stage). To this end, a simple yet useful counting filter idea (Grossi and Luccio, 1989; Jokinen *et al*., 1996) is applied. For a given read and all suffixes in a range, a histogram of pairs of DNA symbols is computed; there are 4^2^ = 16 such pairs useful for our purpose. If the difference between the two histograms, for the read and for the suffix, exceeds 4*i*, it implies that the true distance is greater than *i*.

The other optimization greatly reduces the number of read vs. suffix comparisons in the areas we recognized as difficult. Sometimes we come across a situation that for the current read there are hundreds or thousands of suffixes with an exact match for a considered segment (it happens especially for repetitive segments, e.g., `AAA…A`). It is likely that such a read will be followed, in the sorted bin order, with many reads with the same substring in the considered segment which will imply a lot of read vs. suffix distance calculations. To prevent such bad case, after passing the first read over a difficult area we calculate simple signatures for all the suffixes in the area, in order to compare each of the reads to follow with only a (usually small) subset of suffixes, namely those for which there is still a chance to have the Levenshtein distance within the assumed limit.

We do not present a formal description of this routine. Rather, let us explain it on an example. Let the read length be 100, Hamming distance used, *i* = 3 (i.e., we accept up to 3 mismatches in the current stage and thus partition reads into 4 segments), and let the current segment comprise the range of read symbols [0…24]. We consider a dictionary data structure, whose keys will be short DNA strings of a fixed (small) length and associated values the IDs of suffixes which have the key at a specified position. Assume that the key length is 6. We build *i* + 1 = 4 such dictionaries, and the key positions in suffixes should be far from the current segment. If, for example, the considered suffix points to location *ℓ* in the reference genome 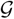, the dictionary keys are: 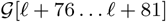, 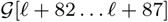, 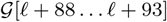, and 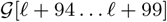.

Now, for the current read *r* we examine the list of suffixes under the key of *r*[76…81]. Those suffixes are potential matches (with up to 3 mismatches) and have to be fully compared versus the read. Then, for the same read, we access the list of suffixes under the key of *r*[82…87] and again perform full comparisons. The procedure is continued for the two remaining keys. Note that if some suffix has at least one mismatch versus read *r* in its *r*[76…81] area, and similarly at least one mismatch against *r*[88…87], *r*[88…93], and *r*[94…99], then in total it has at least 4 mismatches against *r* and thus can safely be rejected.

We noticed that extending this idea to *i*+5 (rather than *i*+1) substrings, and thus *i* + 5 corresponding dictionaries, with the criterion that at least five (rather than one) matches on those substrings trigger a full read vs. suffix comparison, is even more selective in practice and we use this variant in the implementation.

It is interesting and perhaps surprising to note that in difficult areas using those dictionaries allows to reduce the number of suffixes checked against a read by 2–3 orders of magnitude. A similar phenomenon can occur for the Levenshtein distance, but in that case the number of required dictionaries grows (due to “shifts” resulting from indel errors). Despite the fact that the improvement in efficiency for Levenshtein errors is lower than in the Hamming case, it is still significant and makes this idea worthwhile.

Finally, we describe the last, (*k* + 1)th, major stage referred to as *sensitive*. Here we have reads for which there are no matches with up to *k* errors. We basically follow the previous, *k*th, stage here, but the allowed number of errors is set (by default) to as many as *ck*, where *c* is a parameter with default value of 3.0. Yet, the number of segments into which we split those hard reads is still *k* + 1, therefore there is no guarantee to find all matches with up to *ck* errors. In this stage we also discard the reads falling into difficult (in the sense described above) suffix areas, as the slow-down wouldn’t be worth the tiny improvement in accuracy for prospective variant calling.

### Postpreprocesing

The postprocessing stage consists in aggregating mappings of individual reads in order to obtain paired-end mappings. For each bin, related data are loaded from disk and sorted according to the ID, which results in paired reads being adjacent (they differ only in the least significant bit of ID). Let (*r*_1_, *r*_2_) be the analyzed read pair. *G* and *H* indicate the sets of individual mappings of *r*_1_ and *r*_2_, respectively. The aim is to identify mapping pair (*g_i_, h_j_*), *g*i ∈ *G* ∧ *h_j_* ∈ *H*, most appropriate from the point of view of DNA evolution and technical aspects of sequencing. The evaluation of mapping pair is done similarly as in BWA-MEM (Li, 2013): *Q*(*g*, *h*) = *S*(*g*) + *S*(*h*) − *P* (*g, h*), with *S* being an alignment score (linear or affine, depending on the postprocessing step) and *P* indicating a penalty for a deviation from the expected template length. The template length (TLEN), the distance between the leftmost and rightmost mapped base in the pair, is modeled by a normal distribution *N*(*µ*, *σ*). The model is built for each bin separately and its parameters are updated during processing. Function *P* is saturated at arguments *µ* ± 5*σ* not to penalize excessively structural variants which may lead to template lengths much larger than expected. Note that the evaluation function *Q* should not be confused with the mapping quality score (MAPQ) calculated on the basis of the probability of pair alignment being improper (Li *et al*., 2008). As the calculation of MAPQ requires also suboptimal mappings, all pairs found during the procedure are stored in a priority queue *Z* ordered by *Q* value.

At the very beginning of the postprocessing, all individual mappings from *G* and *H* sets are assigned with alignment scores *S* according to the linear gap model. Then, the *Z* queue is filled in the following steps:

1. **Pairing high-quality close mappings**—executed when *G* and *H* are non-empty and at least one of them contains mappings from non-sensitive stage. For each element of *G* and *H* assigned with highest *S* scores we try to find a close mapping (TLEN ∈ [*µ* − 4*σ, µ* +4*σ*]) with opposite orientation in the other set. This is done with the binary search as mappings in *G* and *H* are sorted according to the genomic position.
2. **Rescuing close pairs (Levenshtein)**—if *Z* is empty, for each element of *G* and *H* we try to align mate read with Myers’ bit-parallel algorithm for Levenshtein distance. The scan is performed with less restrictive error thresholds in a wider TLEN interval ([0, *µ* + 5*σ*]) than previously. All identified pairs are inserted to queue *Z*, while individual mappings are added to sets *H* and *G*. All scores *S* are updated using affine gap penalty model.
3. **Rescuing close pairs (clipping and long indels)**—if *Z* is empty, for each element of *G* and *H* we try to align mate read with clipping. This is done by hashing all 20-mers of a query read in a table and matching reference genome in [0, *µ* + 5*σ*] TLEN interval against this table. The procedure additionally detects long indels with flanking regions of length 20 or more perfectly matched to the reference. The collections *Z*, *G*, and *H* are updated during the scan as in (2).
4. **Pairing distant mappings**—we take mappings from *G* and *H* with highest *S* scores, combine them into pairs, and add to queue *Z*.

Note that in the vast majority of cases there exist high-quality mappings for both reads. In these situations, computationally intensive steps (2), and (3) are omitted, resulting in superior execution times.

After filling *Z* queue, the MAPQ scores are assigned to the individual read mappings according to the formula MAPQ = −10 log_10_ (probability of mapping being invalid) saturated at value 60. When the read has only single mapping, it is assigned with the highest MAPQ value. If the single best alignment is accompanied with suboptimal alignments, the MAPQ score is decreased similarly as in BWA-MEM—the misalignment probability is estimated on the basis of the difference between the highest and the second highest score *S*, as well as the number of mappings with the latter. When there are multiple mappings with the highest *S* value, MAPQ decreases vastly. For instance, when there are two equally good alignments, the probability of a mapping being wrong is 1/2 which results in MAPQ = 3. All suboptimal mappings are assigned with MAPQ = 0. After giving MAPQ values to the individual read mappings, the same procedure is applied for pairs, with *S* measure being replaced by *Q*. Finally, reads are assigned with the larger from individual and paired MAPQ scores.

In the output SAM file, a pair mapping with the highest *Q* value is reported. When there are multiple such pairs, one is selected at random. If only one read from pair is mapped successfully, the single-end mapping is reported, while the other read is marked as unmapped. Pairs of reads with no mappings identified during the main processing do not enter preprocessing at all–they are immediately stored as unmapped. To reduce the I/O overhead related to saving large SAM files, Whisper has the ability to compress the output on the fly to gzip format.

## 3 Results

The goal of this section is to experimentally evaluate the accuracy as well as time and memory efficiency of our mapper, Whisper, compared to several leading solutions. Fig. 1 shows the results of variant calling with reference to the human genome NA12878 (Ref HG38) sequenced as part of the Illumina Platinum Genomes (Eberle *et al*., 2017). The reason for this choice is that the National Institute for Standards and Technology (NIST) has released a high-confidence set of variants for that individual as part of the Genome In a Bottle (GIAB) (Zook et al, 2014) initiative. This allows us to consider this set as the “ground truth”. For calling the variants we followed GATK (McKenna *et al*., 2010) Best Practice pipeline (Auwera *et al*., 2013).

**Fig. 1.**
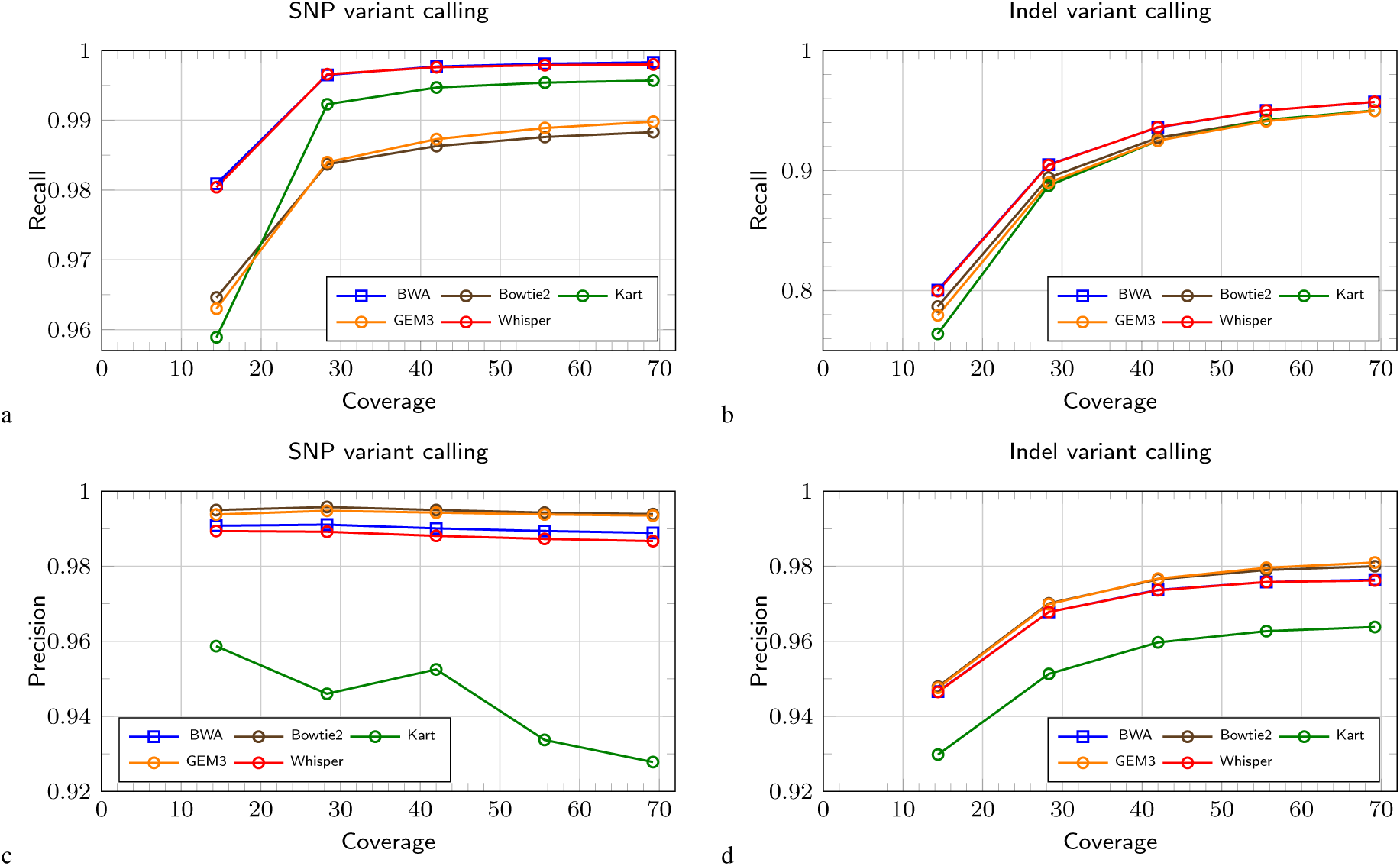
Variant calling in a function of growing coverage, for SNPs (left figures) and indels (right figures). Recall and precision are presented in top and bottom figures, respectively. Detailed results are given in Supplementary Material.

The left figures present VC results for SNPs and the right ones for indels. The top and the bottom row shows the recall and the precision scores, respectively, in function of varying read coverage, up to 69x. The competitors include BWA-MEM (Li, 2013), Bowtie2 (Langmead and Salzberg, 2012), Kart (Lin and Hsu, 2017), and GEM3 (Marco-Sola *et al*., 2012), at their default settings. As expected, for all algorithms, for a given coverage, indels are harder to call than SNPs (all measures are lower for indels). For SNPs, BWA-MEM and Whisper dominate over the others in recall, yet Bowtie2 and GEM3 boast with higher precision. We can say that Bowtie2 and GEM3 try to stay on the safe side and prefer precision over recall. In precision, also BWA-MEM is superior to Whisper. Kart yields relatively high recall but also low precision. Strangely, the precision with SNPs degrades a little (and even noticeably for Kart) for most mappers, a phenomenon which we are unable to explain.

For indels, the trends are basically similar. BWA-MEM and Whisper have practically the same recall and precision curves, and again Bowtie2 and GEM3 are better in precision but worse in recall. Kart is clearly worst in precision, but comparable to GEM or Bowtie2 in recall for high coverages; when the coverage is below 30x it comes last also in recall.

Table 1 focuses on the “typical” coverage of 42x. In mapping speed we have two distinct groups: Whisper, Kart, and GEM3 belong to the faster ones, while BWA-MEM and Bowtie2 are over 6.5 times slower than Whisper. In memory usage, there is an opposite tendency, with BWA-MEM and Bowtie2 being more frugal. Whisper needs the largest amount of RAM memory, yet it easily fits a 32GB (or even 24GB) RAM machine. More detailed timings for Whisper, for varying coverage and for its particular stages, are given in Supplementary Material.

**Table 1.** Results for the real dataset NA12878 (Ref HG38)

| Mapper | Time<br>[h:mm] | Memory<br>[GB] | Unmapped<br>reads [%] | Unmapped<br>bases [%] | SNP |  |  | Indels |  |  |
| --- | --- | --- | --- | --- | --- | --- | --- | --- | --- | --- |
|  |  |  |  |  | Recall | Precision | F1 score | Recall | Precision | F1 score |
| BWA-MEM | 12:55 | 8 | **0.54** | **1.44** | **0.9977** | 0.9901 | **0.9939** | 0.9359 | **0.9737** | 0.9544 |
| Bowtie2 | 13:03 | 4 | 2.02 | 2.02 | 0.9863 | **0.9950** | 0.9906 | 0.9273 | 0.9765 | 0.9513 |
| Kart | 2:27 | 12 | 1.51 | 1.84 | 0.9947 | 0.9525 | 0.9732 | 0.9248 | 0.9597 | 0.9419 |
| Gem3 | 2:36 | 17 | 1.26 | 1.77 | 0.9873 | 0.9943 | 0.9907 | 0.9249 | 0.9767 | 0.9501 |
| Whisper | **1:57** | 18 | 1.51 | 2.24 | 0.9976 | 0.9881 | 0.9928 | **0.9361** | 0.9736 | **0.9545** |

Mapping time, RAM usage, percentage of unmapped reads (resp. bases) and variant calling recall, precision, and F1 scores for real data with 42x coverage. Both SNPs and indels are considered. All tools were run using 12 threads. The best results are in bold.

In variant calling, the results for each mapper are representative to their overall performance, as commented with regard to Fig. 1. Here we also present F1 scores. F1 is a joint measure of recall and precision and can thus summarize the quality of particular algorithms. For SNPs, BWA-MEM is the winner according to F1, with Whisper coming a close second. For indels, Whisper and BWA-MEM are practically equal. For both kinds of variants, Kart is never on a par with the remaining tested tools.

Fig. 2 presents mapping results for simulated reads of length 75, 100, 125, and 150 bp, respectively; the reads were generated with wgsim. 1M read pairs (i.e., 2M reads) were taken for each run. The four bars, in order, stand for unmapped reads, incorrectly mapped reads, and unmapped (resp. incorrectly mapped) reads among those with MAPQ value ≥ 20, i.e., those which may be considered of good quality. The threshold 20 was used in the variant calling experiments on real data. We note that MAPQ values are software-dependent and thus comparing them between different mappers is risky, yet the value of 20 is “understood” similarly by multiple tools. Moreover, this MAPQ threshold is used in the GATK Best Practice pipeline. As expected, the results get consistently better (i.e., bars gets shorter) with longer reads. Among the test set of tools, Kart has somewhat atypical characteristics. Its percentage of incorrectly mapped high-MAPQ reads is often low, but the percentage of unmapped high-MAPQ reads is always the highest; the difference is especially striking for the longest reads. The other four contenders have similar trends, with BWA-MEM and Whisper being noticeably better than GEM3 and Bowtie2.

**Fig. 2.**
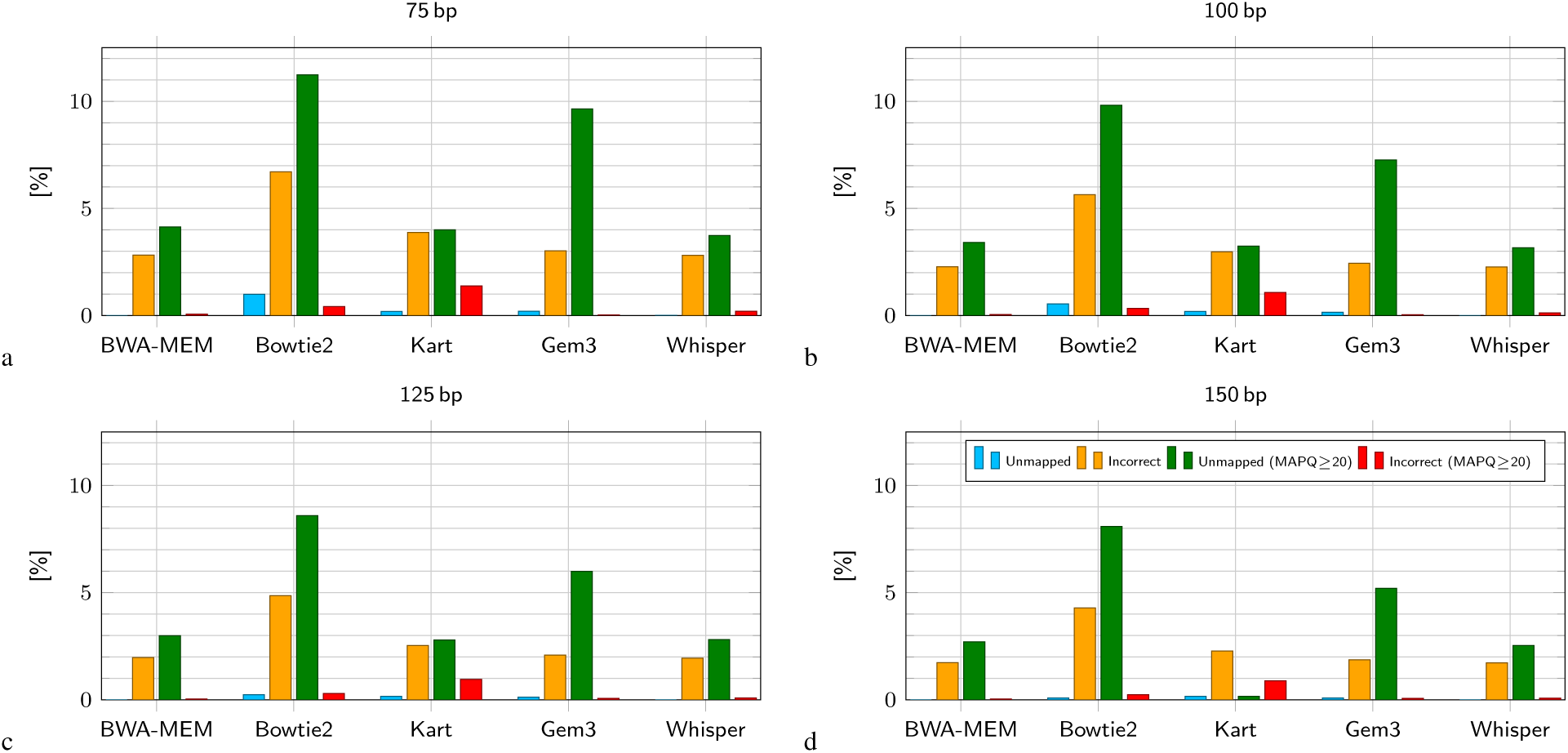
Mapping results for simulated reads of different length. 1M read pairs was taken for each run. Detailed results are given in Supplementary Material.

We believe that comparing accuracies of variant detection is a more appropriate way of benchmarking mappers on real data than using the percentage of mapped reads or even their MAPQ scores estimated by tested tools. This is because not all returned read alignments are relevant and there is no established mapping quality measure in use across all leading mappers. It is variant calling which actually shows if the mappings are relevant. On the other hand, the (still) more widely used statistics of mapped/unmapped reads and their quality (MAPQ) are reasonable for simulated reads.

## 4 Conclusions

We present Whisper, a fast and accurate mapper for NGS reads, handling mismatch and indel errors. It contains a number of novel ideas beneficial both for mapping speed and accuracy. Its general approach of sorting reads and then mapping them in order against a suffix array built for the reference genome (and its reversed complement) is rarely pursued, yet it allows not only high speed, but reasonable memory use, as the suffix arrays may be read in blocks from disk. The processing is performed in many stages, to detect up to *k* errors, based on the pidgeonhole principle. Special care is taken for difficult genome areas, to reduce the number of needed matchings by 2–3 orders of magnitude compared to a more naive approach. Other algorithmic ideas, beneficial for performance, are to use Myers’ bit-parallel edit distance computing routine, the counting filter, and (re)packing DNA symbols in pairs or triples, to reduce the I/O and speed up approximate string matching. We also efficiently utilized the hardware resources: our implementation is highly parallel, using CPU threads and AVX2 or other available SIMD extensions. Temporary data are stored on disk, with care taken to minimize I/O operations.

As a result, Whisper is more than 6.5 times faster than the well-known Bowtie2 and BWA-MEM. It is also by about 30% faster than recently presented GEM3 and Kart, probably the fastest mapping tools nowadays.

Although Whisper essentially handles up to *k* errors, some matches with more (up to 3*k* by default) Levenshtein errors are also detected. More important, however, for high accuracy is careful handling of paired-end reads, which is performed in the postprocessing stage. Mapping pairs are evaluated with a linear or affine alignment score. The distance between the leftmost and rightmost mapped base in the pair is assumed to be modeled by a normal distribution whose parameters are learned. Accuracy evaluation was performed on both real and synthetic reads of varying lengths and coverages. Experiments with real reads via variant detection showed Whisper to be generally at least on a par with Bowtie2 and GEM3 in accuracy and slightly inferior to BWA-MEM (only with SNPs, as the results in indel variant callings are practically identical); we however believe that the almost 7-fold difference in speed poses an attractive tradeoff.

